# Extracellular vesicles from patients with coronary artery disease (CAD) demonstrate enhanced pro-coagulatory activity

**DOI:** 10.1101/2023.09.27.559865

**Authors:** Shin Soyama, Ruihan Zhou, Abigail Whyte, Sharon Mark, Leanne Dymott, Mark Brunton, Roman Fischer, Svenja Hester, Charlie McKenna, Neil Ruparelia, Parveen Yaqoob, Keith Allen-Redpath

**Author notes:** Correspondence: Dr Keith Allen-Redpath Brunel University London, Uxbridge, London, UB8 3PH, United Kingdom.

## Abstract

**Aims:** Extracellular vesicles (EVs) carry unique repertoires of biologically active cargo that hold promise as novel biomarkers and future interventional targets for cardiovascular diseases (CVDs). However, it is unclear as to how the number, location, cellular origin, and size of these EVs within the circulation relate to the development of CVDs such as coronary artery disease (CAD). The current study compared these novel markers in arterial and venous blood samples in individuals undergoing coronary assessment with angiography.. EVs were then characterized from those presenting with and without CAD.

**Methods and Results:** Arterial and venous blood samples were collected from individuals with confirmed CAD following coronary angiography and a matched cohort in whom there was no evidence of disease. EV fractions were isolated from 500µL of platelet free plasma (PFP) by size exclusion chromatography. EVs were analyzed by Nanoparticle Tracking Analysis and flow cytometry to characterize number, size and cellular origin. A thrombin generation assay was used to assess the pro-coagulatory activity of the isolated circulating EVs. Proteomics was used to determine the protein cargo carried within the EVs.

Coagulatory activity of EVs isolated from CAD patients was significantly higher in when compared to controls, although the numbers of EVs did not differ. There were higher numbers of endothelial-derived EVs in arterial blood compared with venous blood. Linear regression models revealed that plasma triacylglycerol concentration and age independently predicted circulating EV numbers in CAD patients. Proteomics revealed several proteins associated with coagulation upregulated in patients with CAD.

**Conclusions:** Although the absolute numbers of EVs in CAD patients were not elevated, EVs in CAD patients had greater pro-coagulant activity, highlighting a potentially important role for EVs in the pathogenesis of CAD.

**Clinical Perspective:** *What is new?:*

- CAD patients displayed a greater EV-induced thrombin generation capability when compated to those with no disease.
- The proteomic profile of arterial-derived EVs in patients with CAD demonstrated several proteins associated with coagulation that may be a risk factor for developing a future event.

*What are the clinical implications?:*

- The identification of EVs in patients with CAD with upregulated coagulation proteins may be a novel biomarker for the development of future events and may identify a high risk patient group that may benefit for more aggressive cardiovascular risk intervention.

## INTRODUCTION

Cardiovascular disease and in particular coronary artery disease (CAD) is the number one cause of death worldwide, affecting approximately 126 million people and increasing global prevalence^1^. Moreover, the complications associated with CAD is considered the principal cause for the development of heart failure in western countries ^2^. The pathological processes driving atherosclerosis underlines the development of CAD. ^3^. Ischaemic complications of CAD results from acute plaque rupture (REF) and the development of in-situ thrombus which is the result of hypercoagulability and hypofibrinolysis^4–5^, platelet hyperreactivity.^6^ In addition, other patient risk factors such as obesity^7^, hypertension^8^, hypercholesterolemia^9^ and hypertriglyceridemia^10^ have been demonstrated to contribute further to the risk of developing CAD and suffering a complication EVs are nanoscale, lipid-bilayer encircled vesicles shed by all cells undergoing activation and/or apoptosis. Although the underlying mechanisms involved are not fully understood, EVs contain features that reflect the state of their parent cell, such as similar biological cargo and cell surface markers, making them a promising biomarker in pathological conditions^11^. Although circulating EVs are found in the blood (and other body fluids) of healthy subjects, they are reported to be elevated in pathological conditions associated with endothelial dysfunction, such as atherosclerosis and thrombosis^12–13^. In recent years, studies have also highlighted how the pathophysiological role of EVs is dependent on their cargo, which constitutes lipids, proteins, and nucleic acids^14–15^. For instance, EVs have been shown to facilitate inflammation and cell death in the vasculature by upregulating pro-inflammatory cytokines (e.g. TNF-α, IL-1β, IL-6, and IL-8) and the expression of adhesion molecules (VCAM-1, ICAM01, and E-selectin) on target cells, leading to the development of atherosclerotic plaque^16–20^. They also express negatively charged Phosphatidylserine (PS) and active Tissue Factor (TF), which are pro-coagulant^21^. Furthermore, in a recent publication from our laboratory, Zhou et al.^22^ reported strong associations between circulating EV numbers and traditional and coagulatory risk markers of CVDs in subjects with moderate CVD risk, shedding light on the potential of EVs as a diagnostic biomarker.

There has been little research to date examining the coagulatory function of circulating EVs in CAD; most research has assessed this based on the quantification of pro-coagulatory EVs, such as PS^+^EVs or TF^+^EVs^23–25^. However, importantly, the most compelling evidence for the pro-coagulatory activity of EVs comes from functional assays^26^, which were employed in the current study. Venous blood has long been used as the primary blood source used in clinical studies given that it is easy to obtain, but importantly there are very few data investigating EVs in arterial blood that may well more accurately reflect underlying pathological processess in the arterial vasculature (e.g. atherosclerosis) and be more clinically relevant both as potential biomarkers and also targets for intervention. We therefore in the present study aimed to determine if there were differences both quantitatively and qualtitatively in indivudals with and without CAD and between arterial and venous blood.

## METHODS

### Subjects and study design

A total of 60 subjects (consisting of 40 patients with confirmed CAD and 20 matched controls) were recruited for the current study. All patients were recruited following invasive or non-invasive coronary angiography in the investigation of CAD. Individuals with confirmed coronary artery disease based on angiography were allocated to the CAD group, and patients with no evidence of CAD were allocated to the control group. Patients presenting with the following criteria were excluded from the study, Acute coronary syndrome in the past 12 months; on P2Y12 inhibitors (including clopidogrel, ticagrelor and prasugrel); On treatment dose anti-coagulation, including warfarin or novel anti-coagulant drugs (Dabigatran, Apixaban, Rivaroxaban, Edoxaban); Metabolic dysfunction; Evidence of alcohol or drug misuse; Inability to give informed consent; Under 18 years old; Pregnancy; Active or recent malignancy (<2 years); Underlying haematological pathologies. All individuals provided written informed consent approved by the Human Research Authority and Health and Care Research Wales (REC 20/NW/0263).

Venous and arterial blood samples (30ml each) were collected from patients undergoing invasive coronary angiography at the Royal Berkshire Hospital, drawn into a vacutainer containing tri-sodium citrate (3.2%; Greiner Bio-One, UK) and processed within 2 hours. Platelet-free plasma (PFP) was prepared by differential centrifugations of 1) 1,500 x g for 15 minutes at room temperature and 2) 13,000 x g for 2 minutes at room temperature at the University of Reading within one hour of collection, and the PFP was subjected to size exclusion chromatography (SEC) and ultra-filtration (UF) steps to isolate and concentrate circulating EVs. Total EV numbers and their subtypes were assessed on the same day using NTA and flow cytometry. Thrombin generation, western blotting, and proteomic analysis were performed on EVs isolated from defrosted PFP samples within 3–6 months of freezing. For EV analysis, a sample size of 34 subjects was deemed sufficient to detect a 10% difference in EV numbers between control and IHD patients with a significance level at 0.05 (two-sided) and a power of 95%, based on a previous study conducted in our research group investigating fish oil supplementation^27^. This calculation assumes that the standard deviation is 2.4 counts/µL (Wu and colleagues observed standard deviations between 1.38 and 2.4 counts/µL for EV counts). 40 subjects were recruited to allow for a 15% dropout rate. Unfortunately, the number of control patients (n=20) was half that of the IHD patients due to the low number of patients undergoing invasive coronary angiography; venous-only blood was collected for this group following computerized tomography (CT) assessment.

### Enumeration and characterization of EVs

#### Nanoparticle Tracking Analysis (NTA)

The size distribution and concentration of circulating EVs were analyzed via NTA using a NanoSight NS300 instrument (Malvern Instruments). Samples were diluted in PBS to achieve a concentration of approximately 1-9×10^8^ particles/ml and injected into the NanoSight sample chamber using a syringe pump. Videos of each diluted sample were analyzed by Nano 3.2 software (Malvern Panalytical).

#### Characterization of EVs by flow cytometry

Flow cytometric analysis was used to characterize circulating EVs based on their number, size and expression of specific markers. Firstly, the limits of detection of the flow cytometer (Canto II Flow Cytometer (BD Biosciences, UK)) and the EV gate were established using standard-size calibration beads (ApogeeMix size range 180-1300nm; Apogee Flow Systems). Events that fell within this gate were classified as being ≤1 µm in size. EVs were then assessed for their ability to bind cell marker-specific monoclonal antibodies such as anti-CD41 (Biocytex, Marseille, France) to detect and count platelet-derived EVs, anti-CD105 to detect endothelial-derived EVs, anti-CD235 to detect red blood cell-derived EVs, anti-human HLA-A,B,C to detect leukocyte derived EVs and Annexin V (ThermoFisher, UK), which binds externalized phosphatidylserine (PS) residues. Data was captured using FACSDiva Software version 6.1.3 and analyzed using FlowJo version 10 (BD).

#### Preparation of concentrated EV suspension by ultrafiltration

The EV fractions from SEC were pooled and concentrated using a VivaspinTM centrifugal concentrator (Fisher Scientific, Leicestershire, UK) at 1000 x g for 10 minutes at room temperature. The protein concentration of the pooled EV suspension was measured using a Nanodrop-1000 spectrophotometer and adjusted to the desired concentration for downstream analysis.

#### Measurement of protein concentration of EVs using NanoDrop

Prior to measurement, the instrument was cleaned with distilled water and blanked using 2 μl of nuclease20 free water. The sample (2 μl) was then loaded onto the pedestal and repeated three times to obtain the average protein concentration.

### Measurement of thrombin generation

Thrombin formation was assessed using a commercially available, plate-based thrombin generation assay (Technothrombin TGA kit, Austria), which evaluates a change in fluorescence as a result of cleavage of a fluorogenic substrate by thrombin over time upon activation of the clotting cascade by tissue factor. Two separate analyses were conducted: (i) determination of the effect of n-3 PUFA supplementation on thrombin generation in PFP from study samples relative to pooled VDP and (ii) determination of the effects of circulating EVs and in vitro-generated PDEVs derived from study samples on thrombin generation in pooled VDP. Before analysis, a thrombin calibration curve was constructed from dilutions of lyophilized Hepes-NaCl-buffer containing 0.5 % bovine serum albumin and ∼1000 nM thrombin in buffer with BSA. A kinetic reading of the plate was initiated by additional of 50 µl of fluorogenic substrate solution containing fluorogenic substrate 1 mM Z-G-G-R-AMC and 15 mM CaCl_2_. Calibration curves were recorded at 37 °C for 10 minutes with 30-second intervals using the plate reader (FlexStation 3, United States) at 360 nm for excitation and 460 nm for emission.

The methods for both approaches were based on using pooled VDP as a negative control to assess thrombin generation specifically resulting from the presence of EVs. For the first approach (i), 40 µl aliquots of either pre-thawed study sample PFP or pooled VDP or pooled PFP were added into the plate. For the second approach, PDEVs (10 µl of EV suspensions at 5 µg/ml final protein concentration) produced from either unstimulated or stimulated platelets (UP-EVs and SP-EVs, respectively) from intervention samples or PBS (negative control) were added to 30 µl VDP. The 5 µg/ml protein concentration was determined through trials as an appropriate concentration for thrombin generation in this assay. This was followed by adding a 10 µl suspension of phospholipid micelles containing recombinant human tissue factor (TF) in Tris-Hepes-NaCl buffer (RCL), provided in the kit. Thrombin formation was initiated by adding 50 µl of fluorogenic substrate solution containing 1 mM Z-G-G-G-AMC and 15 mM CaCl_2_. Plates were immediately read at 37 °C for 1 hour at 1 min intervals using a fluorescence plate reader (FlexStation 3, United States) at excitation and emission wavelengths of 360 and 460 nm, respectively. All samples were measured in duplicate. TGA Evaluation Software detected fluorescence intensity to calculate thrombin generation in samples. The TGA Evaluation Software then analyzed data manually to convert the unit of thrombin generation from RFU to nM and presented as three variables: lag time, time to the peak, peak concentration of thrombin (nM), velocity-index and area under the curve.

### Proteomic analysis

EV suspensions containing 50 µg protein were digested using FASP digestion (Vacacon500, Sartorius, VN01H02 10kDA), and sample volumes were adjusted to 100 µL with 8M urea. The proteins were denatured with 100 µL of 8M urea in 100mM triethylammonium bicarbonate (TEAB) for 30 minutes at room temperature. The samples were then reduced by the addition of Tris (2-carboxyethl)phosphine (TCEP) to a final concentration of 50mM for 30 minutes at room temperature and alkylated by adding carbonyl alkylative amination (CAA) to a final concentration of 50mM for 30 minutes at room temperature in the dark. The filter device containing the mixture was centrifuged at 14,300 x g for 10 minutes at room temperature. Digestion was achieved with the addition of 2.5 µg of trypsin in 200 µL of 50mM TEAB for overnight at 37 °C. Peptides were quenched in two-washes with 200 µL of 0.1% trifluoroacetic (TFA) and 200 µL of 50% acetonitrile (ACN) in 0.1% TFA; each spun through the filter to collect all remaining sample. Collected samples were then dried down in a Speedvac. Dried peptides were reconstituted in 21 µL of 5% formic acid and 5% DMSO and were analyzed by LC-MS/MS on a Fusion-Lumos (Thermo Fischer Scientific, UK). Data was acquired in Data-Independent Acquisition (DIA). A library-free data search was performed in DIA-NN 1.8.1 with the database UPR UPR_Homosapiens_9606_UP000005640_20200803.fasta. False discovery rate (FDR) was set to 0.01, and match between runs (MBR) was enabled. MS intensities were normalized by MaxLFQ (generic label-free technology), and proteins were log(2) transformed.

### Statistical analysis

The Shapiro-Wilk test was applied to assess the normality of the baseline parameters. All variables with non-normal distribution were transformed into logarithmic form for statistical analysis. Unpaired samples t-tests were conducted to compare the means of two groups (IHD vs control), and paired samples t-tests were used to compare the means of two blood types within the IHD subjects (arterial vs venous). Correlations between baseline characteristics and EV parameters were assessed using Pearson’s correlation coefficient or Spearman’s correlation coefficient where appropriate. For those significantly correlated variables, univariate regression analyses were carried out, followed by a multivariate regression analysis in which values of F ≤ 0.05 were included and F ≥ 0.10 were excluded. Finally, to identify independent predictors of EV number, the associated variables were entered into a stepwise regression model. All statistical analyses were carried out using SPSS ver.28.0 (IBM SPSS Statistics, Chicago, IL, USA). Ap-value of 0.05 was considered statistically significant.

## RESULTS

### Patient baseline characteristics

A total of 60 patients were recruited. Final EV analysis was completed for 39 CAD and 20 control subjects. One subject was removed from the analysis due to hemolysation of the blood. The baseline characteristics for CAD patients and controls are shown in Table 1. The CAD patients in the study population were older and had a higher haemoglobin, pro-coagulant parameters (lag time, peak thrombin, velocity index, and ETP) and significantly lower platelet count and TC than the controls. In particular, peak thrombin and ETP were significantly higher in CAD patients when compared to controls (+38.8% peak thrombin and +48% ETP respectively), reflecting greater thrombogenic activity in CAD-EVs. CAD patients were approximately ten years older than controls (64.2 years vs 54.0 years, p<0.05).

**Table 1.** Baseline characteristics of the patients. Data are mean ± SEM. *Denotes a significant difference at *p*<0.05 compared to control. *BMI, body mass index; EDEVs, endothelial-derived extracellular vesicles; eGFR, estimated glomerular filtration rate; ETP, endogenous thrombin potential; HDL-C, high-density lipoprotein cholesterol; IHD, ischaemic heart disease; LDL-C, low-density lipoprotein cholesterol; PDEVs, platelet-derived extracellular vesicles; PS^+^EVs, phosphatidylserine-positive extracellular vesicles; TAG, triacylglycerol; TC, total cholesterol*.

| Characteristics | Controls (n=20) | IHD (n=39) |
| --- | --- | --- |
| Age (years) | 54.0 ± 2.7 | 64.2 ± 1.4* |
| Female:male ratio | 11:9 | 5:34 |
| Height (m) | 1.69 ± 0.02 | 1.74 ± 0.01* |
| Weight (kg) | 86.2 ± 6.7 | 83.5 ± 2.9 |
| BMI (kg/m<sup>2</sup>) | 29.8 ± 1.8 | 27.3 ± 0.7 |
| Haemoglobin (g/L) | 141.1 ± 2.2 | 148.0 ± 2.0* |
| Platelet (thou/μL) | 257.1 ± 9.5 | 230.7 ± 7.3* |
| eGFR (mL/min/1.73m<sup>2</sup>) | 81.1 ± 2.3 | 74.8 ± 1.7 |
| TC (mmol/L) | 4.52 ± 0.21 | 3.89 ± 0.17* |
| TAG (mmol/L) | 1.18 ± 0.13 | 1.07 ± 0.06 |
| HDL-C (mmol/L) | 1.32 ± 0.11 | 1.18 ± 0.07 |
| LDL-C (mmol/L) | 2.67 ± 0.17 | 2.22 ± 0.14 |
| TC/HDL-C ratio | 3.74 ± 0.25 | 3.52 ± 0.15 |
| Circulating EVs (per mL blood) | 2.1 $\times 10^{10} \pm 4.4 \times 10^9$ | 1.7 $\times 10^{10} \pm 2.3 \times 10^9$ |
| EV mean size (nm) | 94.1 ± 2.0 | 91.9 ± 1.4 |
| PS<sup>+</sup>EVs (per mL blood) | 2.6 $\times 10^7 \pm 1.7 \times 10^6$ | 2.7 $\times 10^7 \pm 2.1 \times 10^6$ |
| PDEVs (per mL blood) | 1.4 $\times 10^7 \pm 1.3 \times 10^6$ | 1.3 $\times 10^7 \pm 1.0 \times 10^6$ |
| EDEVs (per mL blood) | 4.0 $\times 10^7 \pm 5.5 \times 10^6$ | 3.8 $\times 10^7 \pm 3.0 \times 10^6$ |
| Lag time (min) | 29.6 ± 1.0 | 25.7 ± 0.6* |
| Time to reach peak (min) | 48.8 ± 1.2 | 47.2 ± 1.0 |
| Peak thrombin (nM) | 58.2 ± 5.0 | 80.8 ± 4.4* |
| Velocity index | 3.1 ± 0.3 | 3.9 ± 0.3* |
| ETP (nM thrombin x min) | 888.7 ± 116.6 | 1315.7 ± 88.7* |

### Greater thrombin generation associated with EVs from venous blood of CAD patients compared to controls

The addition of EVs from venous blood to VDP significantly enhanced thrombin generation (Figure 1). Circulating EVs from CAD patients had a greater capacity to support thrombin generation when compared to controls, with a significantly shortened lag time (p<0.001), a significantly increased peak thrombin concentration (p<0.01), a greater velocity index (p<0.05) and a greater AUC (p<0.05) (Figure 1). Analysis of venous-derived EVs from CAD and control patients revealed no differences in EV numbers, particle size or size distribution (Supplementary Figure 1). In addition, no significant differences were detected in the numbers of PS^+^EVs, PDEVs, or EDEVs (Supplementary Figure 2).

**Figure 1.**
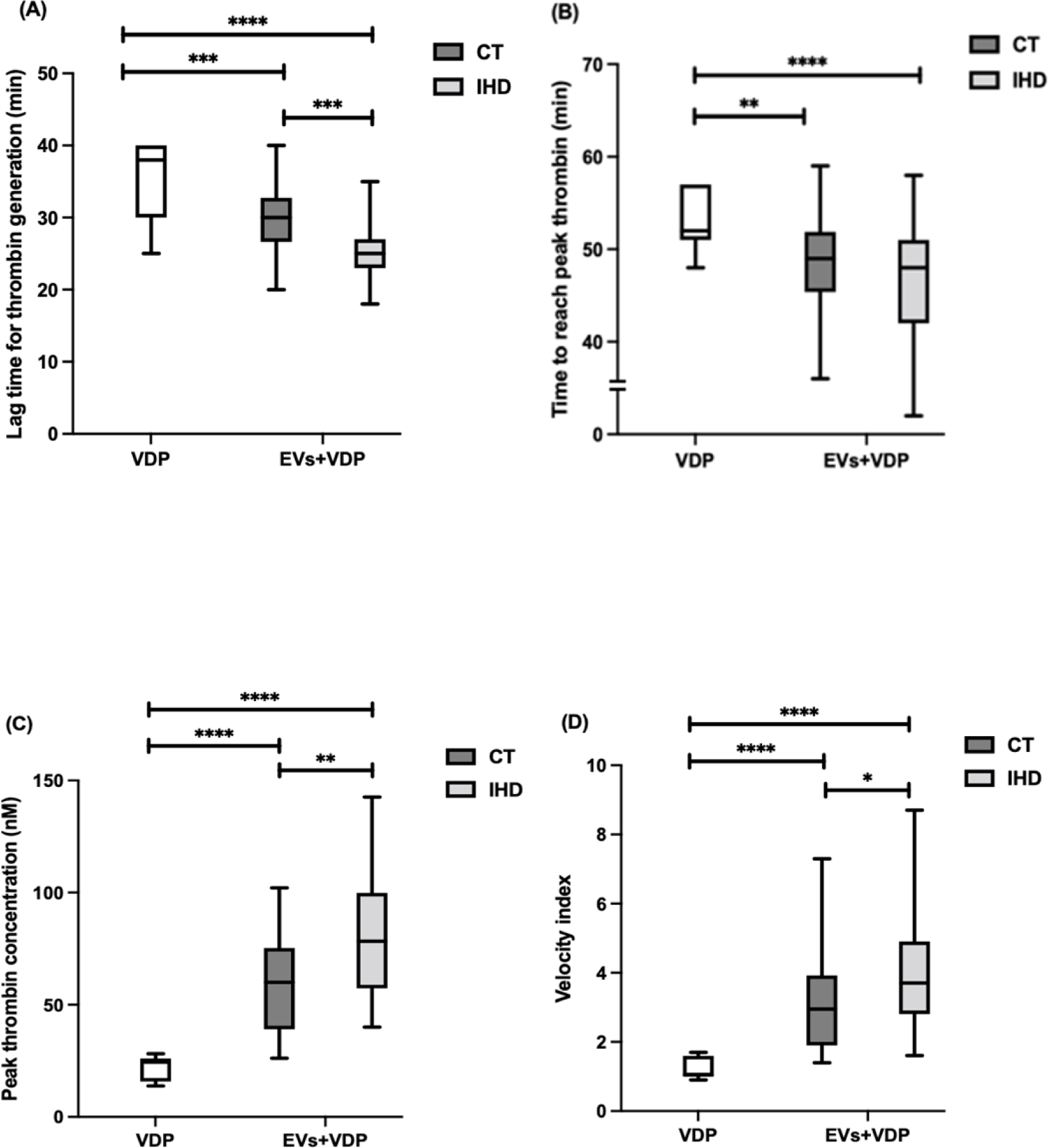
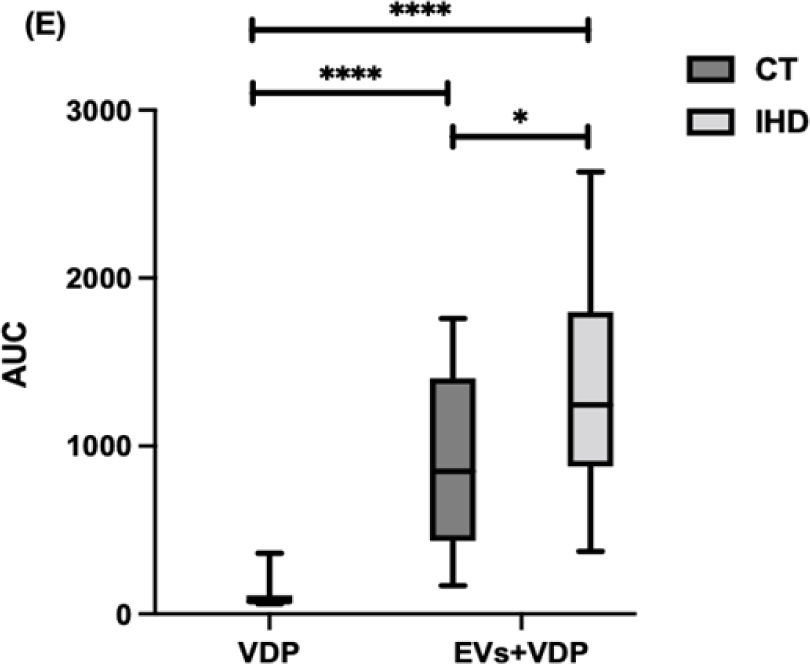
EV-dependent thrombin generation in IHD and control patients (venous blood). Data are median ± 75^th^ and 25^th^ percentiles (boxes) and maximum and minimum values (lines) for n= 39 IHD samples and n=20 control samples. **(A)** Lag time required for thrombin generation (*p*<0.001). **(B)** Time required to reach peak thrombin (*p*=0.42). **(C)** Peak thrombin concentration (*p*<0.01). **(D)** velocity index (*p*<0.05). **(E)** AUC, endogenous thrombin potential (*p*<0.05). *Significant differences between the groups are denoted as *p<0.05, **p<0.01, ***p<0.001, ****p<0.0001. AUC, area under curve; CT, control; EVs, extracellular vesicles; IHD, ischaemic heart disease; VDP, vesicle-depleted plasma*.

### Differential location of EVs did not significantly affect thrombin generation *ex vivo*

Comparison of EV numbers and function in arterial and venous blood was analysed in patients with confirmed CAD.No differences in thrombin generation were noted in EVs from arterial vs venous and was also true for lag time for thrombin generation, peak thrombin concentration, time to reach peak thrombin concentration, velocity index or AUC, although both types of EVs significantly enhanced thrombin generation when added to VDP (Figure 2). There were no significant differences in numbers, size or size distribution of total circulating EVs between arterial and venous blood (Supplementary Figure 3). However, arterial blood had a significantly higher number of EDEVs than venous blood (Supplementary Figure 4). There were no significant differences in the numbers of PS+EV and PDEV between the two types of blood (Supplementary Figure 4).

**Figure 2.**
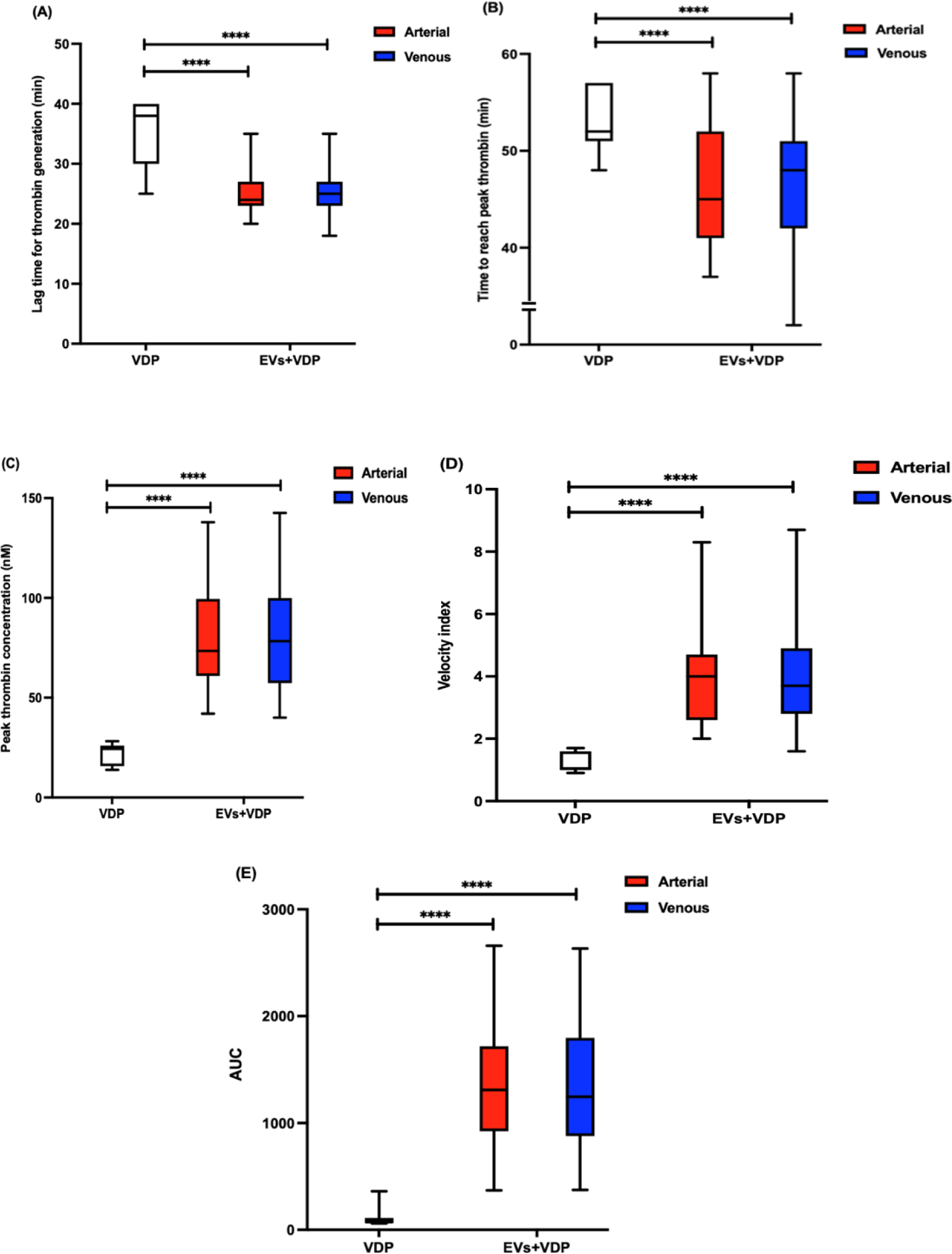
Thrombin generation capacity resulting from the addition of EVs from arterial or venous blood from IHD patients to VDP. Data are median ± 75^th^ and 25^th^ percentiles (boxes) and maximum and minimum values (lines) (n=39). **(A)** Lag time required for thrombin generation (*p*=0.57). **(B)** Time required to reach peak thrombin (*p*=0.37). **(C)** Peak thrombin concentration (*p*=0.85). **(D)** Velocity index (*p*=0.89). **(E)** AUC, endogenous thrombin potential (*p*=0.59). *\*\*\*\*denotes a significant difference at p<0.001. AUC, area under curve; CT, control; EVs, extracellular vesicles; IHD, ischaemic heart disease; VDP, vesicle-depleted plasma*.

### Comparative proteomics and enrichment analysis of EVs highlights key differences between CAD versus control patients

In arterial EVs, a total of 567 proteins were identified, of which 370 proteins (65.3%) were found in both CAD and control EVs, 196 proteins (34.6%) were exclusive to CAD EVs, and 1 protein (0.2%) was exclusive to control EVs (Figure 3A). Bioinformatic analysis of the identified proteins was performed to identify clusters of differentially upregulated proteins associated with specific functional/biological pathways. Proteins involved in neutrophil degranulation, complement and the coagulation cascade and haemostasis appeared to be upregulated in CAD EVs (Figure 3B).

**Figure 3.**
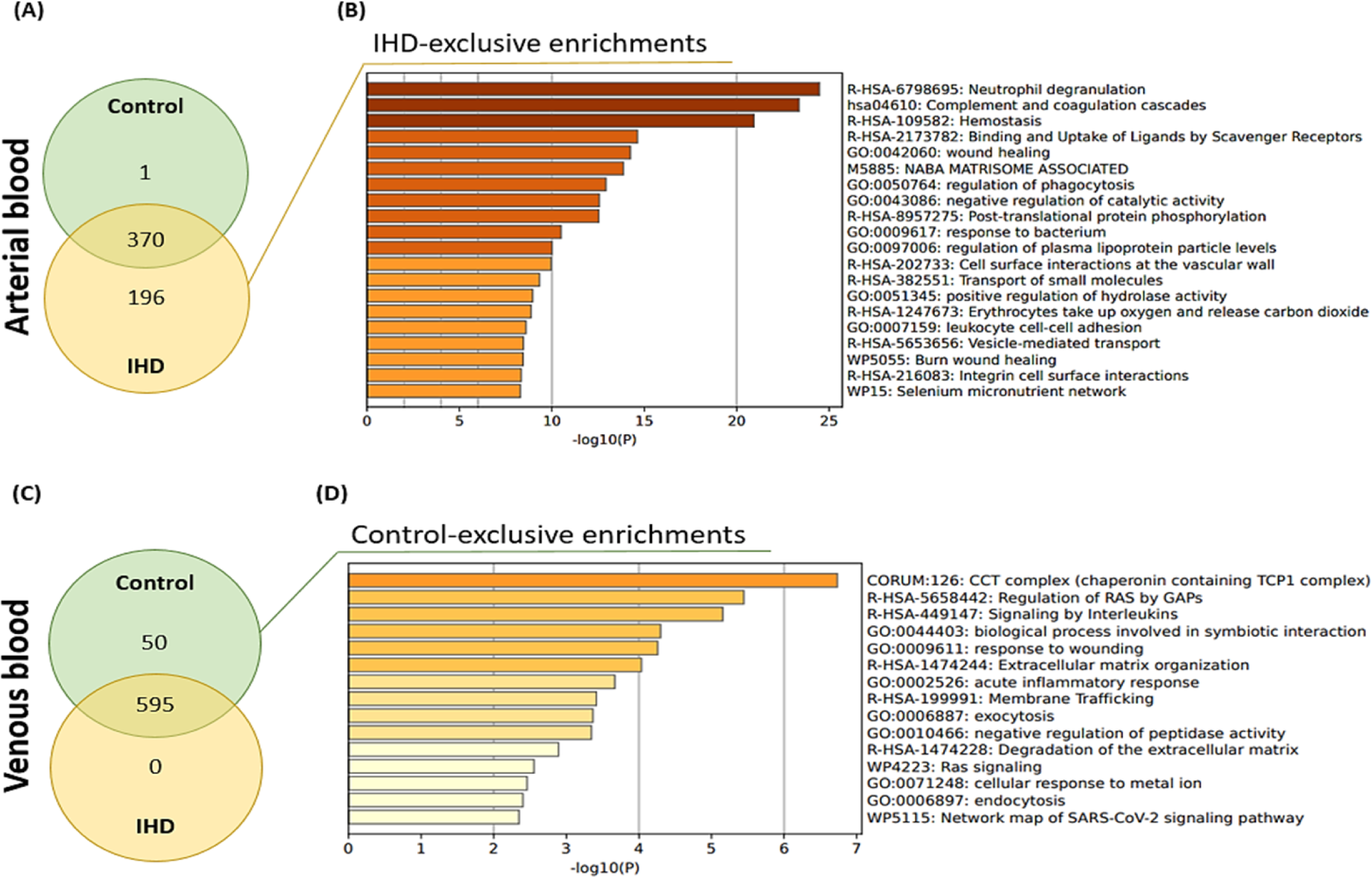
Proteomic profiling and enrichment analysis of EVs from IHD and control patients derived from arterial or venous blood (n=1). **(A)** Venn diagram illustrating the number of proteins identified in EVs derived from arterial blood and **(B)** downstream enrichment analysis of IHD-exclusive proteins. **(C)** Venn diagram illustrating the number of proteins found in EVs from venous blood and **(D)** downstream enrichment analysis of control-exclusive proteins. *GAPs, GTPase activating proteins; IHD, ischaemic heart disease*.

In venous EVs, a total of 645 proteins were identified, of which 595 proteins (92.2%) were present in both CAD and control EVs and 50 proteins (7.8%) were expressed only in control EVs (Figure 3C). There were no proteins exclusively expressed in CAD EVs. Of those proteins expressed exclusively in control EVs, chaperonin containing TCP1 complex, regulation of RAS by GAPs, and interleukin signalling was most notable (Figure 3D).

### Plasma TAG and TC/HDL-C ratio are positively correlated with circulating EV numbers in the venous blood of IHD and control patients, and plasma TAG is positively correlated with circulating EV numbers and EDEVs in the arterial blood of CAD patients

In CAD patients, total EV numbers were positively associated with plasma TAG concentration and TC/HDL-C ratio, and negatively correlated with age and HDL-C concentration (Supplementary Table 1). Platelet counts were positively associated with thrombin generation (AUC and peak thrombin concentration). In controls, total EV counts were positively associated with plasma TAG concentration, TC/HDL-C ratio and thrombin generation (AUC) and negatively correlated with age (Supplementary Table 1). In arterial blood, total circulating EV numbers were positively associated with plasma TAG concentration and negatively associated with HDL-C in CAD patients (Supplementary Table 2). Arterial blood showed numbers of EDEVs were positively associated with plasma TAG concentration, but there was no association for PS+EVs or PDEVs, which was in contrast to venous blood, where PS+EV and PDEV numbers were associated with plasma TAG concentrations whereas EDEV numbers were not (Supplementary Table 2).

Univariate regression analysis showed that in CAD patients, total EV counts were associated with age, plasma TAG concentration, plasma HDL-C concentration, and TC/HDL ratio and in controls they were associated with age, TAG, TC/HDL ratio. However, after multivariate linear regression analysis, EV number remained associated only with age and plasma TAG concentration for CAD patients and plasma TAG concentration for controls (Supplementary table 3). Univariate and multivariate regression analysis confirmed a strong association between total EV numbers and plasma TAG concentration in arterial blood, similar to venous blood (Supplementary table 4).

## DISCUSSION

This study demonstrated for the first time that there are no differences in the number or thrombogenicity of EVs between arterial and venous blood. However, EVs from CAD patients had a significantly greater capacity for thrombin generation than controls.In addition, a strong predictive relationship between circulating EV numbers and plasma TAG concentration was discovered, reinforcing previous observations in healthy subjects^22,27,52^. Furthermore, pilot work using untargeted proteomics suggested some enrichment of coagulation-related proteins in the arterial EVs of CAD patients and some differences in the protein profile of arterial vs venous EVs.

Circulating EVs have been shown to associate with disease severity, and there was an expectation that EV numbers would increase due to the presence of coronary artery disease ^28^. However, there were no significant differences in the number or phenotype of EVs between CAD and control patients. Clinical and observational studies reporting on associations between CAD and the number of EVs are conflicting, some reporting elevated numbers of CD144^+^, CD31^+^/PS^+^ EDEVs and CD41^+^ PDEVs in CAD patients ^29–31^, whilst others report lower CD31^+^/42b^-^ EDEVs in CAD patients compared to controls^32^. This discrepancy may result from the diverse use of ligands for EV labelling, varied baseline characteristics, and other confounding factors (e.g. medications).Since there is no consensus on a specific universal ligand to label EVs, studies have inevitably examined different EV populations, making comparing data across other laboratories challenging. There has previously been no direct comparison of EVs from arterial and venous blood, although there is a limited amount of data comparing the two blood types more broadly. In 28 CAD patients, Kafian and co-workers (2011) reported similar platelet aggregation between arterial and venous blood using a Multiple Electrode Aggregometry device^33^. Separately, blood counts, including platelets, were also reported to be similar between venous and arterial blood of 12 healthy subjects^34^.

Moreover, in an animal study conducted by Doering et al. (2014), thromboelastrography revealed that overall clot formation did not differ between arterial and venous blood in swine, which may explain the lack of coagulatory difference observed in the current study between the two types of blood^35^. The present study, however, demonstrated that arterial blood of IHD patients expressed higher levels of EDEVs compared to venous blood, possibly indicating stronger activation of endothelial cells in the arteries. Comparative proteomics revealed some differences in the protein content of EVs from arterial and venous blood, but these were quite limited and, as an exploratory study, should be considered preliminary data only. It is also worth noting that focusing only on the protein profile of EVs provides a limited view of their potential bioactivity; recent advances in targeted lipidomics are revealing lipids as key bioactive mediators^36^, which should be explored in the context of the role of EVs in CAD which is beyond the scope of this manuscript. The results of our study would therefore support the further use of arterial blood EVs when investigating their role in the pathogenesis of CAD and also their use as a biomarker or target for intervention.

Historically, links between EVs and thrombogenicity have been limited to examining numbers of ‘pro-coagulant’ EVs, such as PS^+^EVs^23,25^, based on the fact that surface exposure of PS confers pro-coagulant activity onto EVs^37^. However, the current study showed that EVs from CAD patients exhibited significantly greater coagulatory activity despite the numbers of PS^+^EVs being similar to controls. Functional assays such as TGA are an effective means of directly assessing pro-coagulatory activity by EVs^26,38^. The current study employed this technique to provide clear evidence that circulating EVs triggered TF-dependent thrombin generation in VDP, implicating the contribution of EVs to coagulation via the TF coagulation pathway^37,39^. Several studies have demonstrated a pro-coagulative role for TF^+^EVs in vascular disease, including CAD^40^, myocardial infarction^24^, and venous thromboembolism^41^. TF^+^EVs are associated with atherosclerotic plaque burden in subjects with high CVD risk, suggesting that TF^+^EV are functional and may regulate CAD progression^40^. Although the present study did not identify enrichment of pro-coagulative proteins in the venous EVs of CAD patients, proteomic analysis identified 196 unique proteins in EVs from arterial blood, which did reveal CAD-specific enrichment of several pro-coagulative proteins, including F13A1 (coagulation factor XIII A subunit), F13B (coagulation factor XIII B subunit), FGA (Fibrinogen alpha subunit) and FGB (fibrinogen beta subunit), whilst the control EVs showed none of these. The underlying mechanisms involved in the greater coagulatory activity of EVs in CAD are still unknown; nevertheless, investigation of enhanced thrombin generation in circulating EVs from the arterial blood that is likely to be more likely to represented the underlying biological processes of the atherosclerosis in CAD patients is highly novel and provides insight into their potential clinical relevance.

Elevated plasma TAG has long been recognized to increase the risk for CVDs^42^. The results from the current study demonstrate that plasma TAG concentration is a strong independent predictor of total circulating EV numbers, in agreement with previous studies^22,27^. In addition, there was an inverse relationship between circulating EV numbers and age in all subjects, and in CAD, age and TAG independently predict total EV numbers. However, in CAD patients, but not in controls, the association with plasma TAG was also evident in PS^+^EVs and PDEVs. The literature has limited evidence on how increased plasma TAG concentrations may directly influence EV release. High concentrations of plasma TAG have been reported to be associated with endothelial and vascular dysfunction^43–44^ and upregulation of soluble adhesion molecules, such as VCAM-1, ICAM-1, and E-selectin^45^, which are expressed on the surface of EVs^46–47^. Platelet reactivity and platelet markers, such as expression of P-selectin, are also enhanced by increased concentrations of plasma TAG^48–50^.

In the present study, an increase in EV number paralleled an increase in TC/HDL-C ratio for all subjects, consistent with reports that circulating EV number is negatively associated with HDL-C in subjects displaying CVD risk factors. However, this association was not evident in subjects without risk factors^51^. Another study also demonstrated significant associations of increased EDEV number with greater TC/HDL-C ratio and lower HDL-C in 844 healthy subjects^52^. Moreover, inducing a hypercholesterolaemic state in monocytes *in vitro* results in the externalization of PS on the cell membrane and releases PS-bearing EVs^53^. Studies have also demonstrated attenuated production of EVs following the administration of cholesterol-depleting statins in hyperlipidaemic patients, including atorvastatin^54^ and pravastatin^55^, which further highlights the direct association between EV numbers and cholesterol observed in the present study. What is not clear is why the strongest relationship between plasma lipids and EV number is with plasma TAG rather than cholesterol. The predictive relationship for plasma TAG is strong and has been demonstrated consistently by our research group, but the underlying mechanisms remain to be clarified.

Growing evidence has portrayed a strong association between circulating EV numbers and clinical parameters, such as plasma TAG. For example, in Sinning and co-workers’ (2011) study, IHD patients (n=196) had higher average plasma TAG concentration compared to the patients from the current study (1.50 mmol/L vs 1.07 mmol/L, respectively), which may have contributed to the significant higher EV abundance observed in that study. Moreover, studies on clinical patients cannot exclude those on prescribed medication, which is difficult to control and may impact EVs in currently uncharacterized ways. Although there were no significant associations between aspirin/statin intake and total EV numbers in the current study, there is, as yet, no consensus about the potential effect of statins on the number of EVs, as some studies have demonstrated decreased EV numbers^56–57^, whilst others have demonstrated increased EV numbers^58^ or no effect^59^. Concerning aspirin, Connor et al. (2016) demonstrated that incubation of platelets with aspirin inhibited EV formation^60^, and daily aspirin administration for 10 days in diabetic patients reduced the number of EVs of various origins^61^. Although subjects with recent ACS (<12 months) were excluded in the present study, we included patients with (or without) CAD and ischaemia were not evaluated.

## CONCLUSION

This study demonstrated markedly greater coagulatory activity of EVs in CAD patients compared to asymptomatic controls, with no alteration in the total circulating EVs or EV subtypes. Although the precise mechanisms are unclear, they may coincide with a strong independent association between total EV number and plasma TAG concentration and elevated levels of EDEVs, as observed in CAD patients. Preliminary shotgun proteomics demonstrated that EVs from CAD and control patients exhibited differences in protein composition, which could provide new insights into the role of EVs in IHD in the future. EVs may putatively serve as a promising target for identifying CAD-related biomarkers.

## Sources of funding

This project was funded by the Joint Academic Board between the Royal Berkshire NHS Foundation Trust and the University of Reading.

## Disclosures

None.

## Supplemental Material

Tables S1–S4

Figure S1–S4

## Notes

### Competing Interest Statement

The authors have declared no competing interest.

